# Endothelial AMBRA1 loss contributes to vascular dysregulation facilitating metastatic progression in early-stage melanoma

**DOI:** 10.64898/2026.09.29.755318

**Authors:** Laura Walker, Roisin Stout, Martina Di Rienzo, Katie Best, Sofia Bisegna, Luca Occhigrossi, Ioana Cosgarea, Philip Sloan, Grant Richardson, Raffaele Strippoli, Mauro Piacentini, Gian Maria Fimia, Penny E Lovat, Jane Armstrong

**Author notes:** Corresponding author: Professor Jane Armstrong John Dawson Drug Discovery and Development Research Institute Faculty of Health Sciences and Wellbeing University of Sunderland Sciences Complex City Campus SUNDERLAND SR1 3SD, UK. Joint senior authors (Penny E Lovat, Jane Armstrong).

## Abstract

Validated biomarkers for identifying patients with early-stage melanomas at high risk of metastasis are limited. We have previously shown that loss of Activating Molecule in Beclin-1-Regulated Autophagy (AMBRA1) and loricrin in the epidermal microenvironment is associated with tumour recurrence. Here we show that endothelial AMBRA1 deficiency contributes to vascular dysfunction and melanoma progression. Melanoma-conditioned media or exposure to TGFβ1-3 ligands induced post-transcriptional loss of AMBRA1 in endothelial cells. Transcriptomic profiling in both in vitro and human melanoma single-cell RNA sequencing datasets showed activation of cell-cycle and metabolic programmes alongside suppression of endothelial junction, adhesion and immune-supportive pathways in AMBRA1-deficient endothelial cells. Functionally, AMBRA1 loss increased endothelial cell proliferation, accelerated early tubulogenesis, impaired three-dimensional spheroid adhesion, stabilised endothelial-to-mesenchymal transition-associated transcription factors Slug and Snail, and reduced Claudin-5, VE-cadherin and N-cadherin, indicative of endothelial plasticity and dysfunction. Furthermore, loss of AMBRA1 in a subset of intratumoural and peritumoural blood and lymphatic vessels correlated significantly with metastatic progression in a cohort of 121 non-ulcerated primary AJCC stage I/II melanomas classified as AMBLor at-risk. Collectively, these findings position endothelial AMBRA1 as a regulator of vascular plasticity, dysfunction, and immune-supportive endothelial function in early-stage melanomas, and a marker of a permissive microenvironment associated with metastasis.

## Introduction

Cutaneous melanoma is a highly aggressive form of skin cancer, with rising global incidence and mortality rates, presenting a considerable challenge to public health (1). Despite advances in targeted and immunotherapy, tumour recurrence remains a clinically significant challenge (2,3). Current American Joint Committee on Cancer (AJCC) staging criteria are unable to identify patients with early-stage (AJCC stage I/II) melanoma at risk of progression and metastatic relapse (4,5). Recurrence rates range from approximately 10-20% in stage IA/IB to 37-59% in stage IIA/IIB/IIC disease (6), highlighting the urgent unmet clinical need for improved stratification of biologically heterogenous early-stage disease to more accurately identify patients at greatest risk of recurrence and disease progression.

Increasing evidence suggests that tumour progression and patient response to therapy are influenced not only by tumour-intrinsic features, but also by the surrounding tumour microenvironment (TME). Endothelial dysfunction, impaired immune-cell trafficking and immune-excluded microenvironments are important determinants of disease progression and immune responsiveness (7–10). As endothelial cells regulate vascular integrity, angiogenic organisation and leukocyte extravasation, endothelial dysfunction may represent early upstream events that promote tumour dissemination and immune evasion (8,9). Defining mechanisms driving vascular dysfunction in early-stage melanoma may reveal clinically actionable determinants of disease progression and recurrence.

Activating Molecule in Beclin-1-Regulated Autophagy (AMBRA1) is a multifunctional scaffold protein that regulates autophagy, cellular homeostasis and cell-cycle progression (11–15). Beyond its role in autophagy, AMBRA1 suppresses cell proliferation through regulation of key cell-cycle drivers, most notably c-Myc and cyclin D, and its loss promotes tumorigenesis across multiple cancer types (14,15). In melanoma mouse models, tumour-specific AMBRA1 loss accelerates tumour growth and invasion through activation of epithelial-to-mesenchymal-like transcriptional programmes, modulation of anti-tumour immunity, and hyperactivation of focal adhesion kinase (FAK) signalling (16–18). Although these findings highlight the tumour-suppressive role of AMBRA1 in melanoma progression, its role within the broader TME remains unclear.

Transforming growth factor beta (TGFβ) signalling can influence the melanoma microenvironment by regulating the proliferation, differentiation, and phenotypic plasticity of surrounding cells in a context-dependent manner (19,20), and also plays a crucial role in endothelial cell function, primarily through induction of endothelial-to-mesenchymal transition (EndoMT) (21,22), modulation of angiogenesis (23), and maintenance of vascular barrier function (24). Previously, we have shown that the combined loss of AMBRA1 and the terminal differentiation marker loricrin in the epidermis overlying non-ulcerated AJCC stage I/II melanomas is associated with an increased risk of tumour recurrence and metastasis (25), where epidermal AMBRA1 loss is driven by TGFβ2 signalling (26).

Here we show that sustained TGFβ signalling drives post-transcriptional endothelial AMBRA1 loss, leading to endothelial dysfunction, characterised by enhanced proliferation, impaired junctional integrity, aberrant vascular remodelling and suppression of inflammatory and leukocyte-recruitment programmes. Furthermore, we show AMBRA1 loss within intratumoural and peritumoural blood and lymphatic vessels is associated with increased risk of metastasis in AJCC stage I/II melanomas. Together, these findings identify endothelial AMBRA1 as a regulator of vascular integrity, and a potential biomarker for high-risk early-stage melanomas, linking tumour-derived TGFβ-driven signalling to endothelial dysfunction and metastatic progression.

## Methods Patient cohort

A retrospective cohort (*n* = 121, Supplementary Table 1) of non-ulcerated AJCC stage I/II defined as AMBLor ‘at-risk’ melanomas (AJCCv8) was collated from three independent diagnostic specimen collections (Melbourne, Australia; Buffalo, USA; Belfast, Northern Ireland), as previously described (25). All cohorts were accessed following full ethical approval through the Newcastle University Dermatology Biobank (REC REF 24/NE/0014) and in accordance with recognised ethical guidelines of the Declaration of Helsinki.

### Semi-quantitative analysis and scoring of endothelial AMBRA1 expression

Automated IHC staining and AMBRA1/loricrin scoring were performed as previously described (25). High-resolution digital images of AMBRA1 stained sections were captured using an Aperio AT2 slide scanner. Areas of each tissue section were labelled as intratumoural, peritumoural (within 500 µm of the tumour boundary) and marginal (dermis, excluding the hypodermis), and the inner layer of vascular cells of each blood or lymphatic vessel was manually annotated across each area based on morphology and characteristic endothelial architecture. Tumour samples with a small margin area (< 10 vessels) were excluded. AMBRA1 expression levels (the number of pixels above a set threshold intensity) within each vessel were determined relative to vessel (endothelium) area using Leica Aperio ImageScope (v12.4.6). The average AMBRA1 expression score across all vessels within each tissue section was determined, from which, the proportion of intratumoural and peritumoural vessels with low AMBRA1 (< 0.45 x average AMBRA1 score) was calculated. A receiver operating characteristic curve was used to determine the optimal threshold for definition of ‘low’ AMBRA1 expression to discriminate between non-metastatic and metastatic tumours in a sub-cohort (*n* = 47). Based on an effect size of 0.892 (observed in the sub-cohort), power calculations (G*Power 3.1.9.2) indicated a cohort of 90 patients was required to obtain 90% power to detect a statistically significant difference at the *P* = 0.05 level in a Mann-Whitney test. Tumour boundary annotation accuracy was confirmed by the study pathologist (PS), and both tissue section annotations and AMBRA1 analysis were performed blind to clinical outcome.

### Cell culture

Human microvascular endothelial cells-1 (HMEC-1, ATCC® CRL-3243™, RRID:CVCL_0307) and primary human dermal blood endothelial cells (HDBEC; PromoCell C-12225) cells were cultured in Endothelial Cell Growth Medium MV (PromoCell C-22020), primary human umbilical vein endothelial cells (HUVEC; PromoCell C-12200) cells were cultured in Endothelial Cell Growth Medium (PromoCell C-22010), CHL-1 (ATCC® CRL-3619™, RRID:CVCL_1122) and A-375 (ATCC® CRL-1619™, RRID:CVCL_0132) cells were cultured in Dulbecco’s Modified Eagle’s Medium (DMEM; Sigma-Aldrich, D6546) at 37 °C in an atmosphere of 5% CO_2_. Primary endothelial cells (HUVEC and HDBEC) were certified mycoplasma-free by the manufacturer and used for experiments only up to passage number 6. HMEC-1 cells were routinely tested for mycoplasma contamination in-house every month.

### Drug treatments and melanoma-conditioned media preparation

Human TGFβ1, TGFβ2 or TGFβ3 recombinant protein (PeproTech®, 17881453, 17860643 and 17831023, respectively) were dissolved in filter-sterilised 10 mM citric acid pH 3.0/0.1% (w/v) BSA in diH_2_O (TGFβ1) or 0.1% (w/v) BSA in diH_2_O (TGFβ2 and TGFβ3) as per the manufacturer’s instructions. Cells were treated every 24 hrs for either 24 or 72 hrs to activate signalling pathways. MG132 (Sigma-Aldrich, 474790) was dissolved in dimethyl sulfoxide (DMSO) (Sigma-Aldrich, D4540) at 10 mM as a stock solution. An equal volume of vehicle was used as a control in all experiments. For melanoma-conditioned media experiments, HMEC-1 cells were seeded onto 6-well plates (3 x10^5^ cells per well) in Endothelial Cell Growth Medium for 24 hrs and then replaced with conditioned media from CHL-1 or A375 cells for 24 and 72 hrs. Conditioned media from CHL-1 or A375 cells were collected after 48 hrs of incubation.

### siRNA transfection

siRNA for AMBRA1 (ON-TARGETplus Human AMBRA1 siRNA SMARTpool, Dharmacon, L-029987-01), or AMBRA1 siRNA #1 and #2 (ON-TARGET custom siRNA, Dharmacon, CTM-1113689 and CTM-1113698), DDB1 siRNA #1 and #2 (ON-TARGETplus Human DDB1 siRNA, Dharmacon, J-012890-06-0010 and J-012890-07-0010), or non-targeting control (ON-TARGETplus Non-targeting, Dharmacon, D-001810-03-05; for ON-TARGET custom AMBRA1 siRNAs, siGENOME Non-targeting siRNA #2, Dharmacon, D-001210-02-50) were transfected into cells using Lipofectamine™ RNAiMAX Transfection Reagent (Invitrogen) according to the manufacturer’s handbook. Cells were incubated for 24-96 hrs at 37 °C prior to cell harvest for gene knockdown studies. For proliferation assays, cells were re-seeded 48 hrs after siRNA treatment, and cell confluency quantified over time using the IncuCyte™ live-cell analysis system.

### RNA processing and quantitative reverse transcription PCR (RT-qPCR)

RNA was isolated from cells using the ReliaPrep™ RNA Miniprep System (Promega) or TRIzol reagent (Invitrogen) according to the manufacturer’s instructions. RNA was converted to cDNA using the High-Capacity cDNA Reverse Transcription kit (ThermoFisher Scientific) or Reverse Transcription Kit (Promega, A3500) according to the manufacturer’s instructions. RT-qPCR was performed using PowerTrack™ SYBR Green Master Mix (ThermoFisher Scientific) or Maxima SYBR Green/ROX Master Mix (ThermoFisher Scientific) and custom primers (Sigma-Aldrich; Supplementary Table 2), using a Rotor-Gene Q2Plex (Qiagen) thermocycler or QuantStudio 3 Real-Time PCR System (ThermoFisher Scientific). The comparative *C_t_* method (2^−ΔΔ^*C_t_*) was used for relative quantification of gene expression.

### RNA sequencing and Gene Set Enrichment Analysis

Total cellular RNA was sequenced (Azenta) generating >20 million reads per sample. Sequencing reads were quality controlled, trimmed and aligned to the human genome (GENCODE v43) using STAR. Gene-level read counts, differential gene expression analysis and Principal Component Analysis (PCA) were performed using DESeq2 in R. Gene set enrichment analysis (GSEA) was performed on a ranked gene list derived from differential expression analysis using the Broad Institute GSEA tool and ClusterProfiler, interrogating Hallmark, KEGG and GO gene sets. Pathways with a false discovery rate (FDR) < 25% and a *P* value < 0.05 were considered significant.

### Curation and analysis of an EndoMT gene signature

A bespoke EndoMT-focused gene signature was assembled based on review of published literature. Genes were manually curated into functional groups reflecting endothelial identity, transitional states, EndoMT inducers, transcriptional regulators, and mesenchymal effector programmes (Supplementary Table 4). This signature was used for visualisation of differentially expressed genes (DEGs) and as a custom gene signature for GSEA.

### TCGA data analysis

TCGA-SKCM RNA-sequencing transcript per million (TPM) data were obtained using the R package TCGAbiolinks (27,28) and analysed at the gene-symbol level. Expression values were log_2_-transformed and compared between primary and metastatic tumours using Wilcoxon rank-sum tests with multiple-testing correction.

### Analysis of GSE72056

Publicly available single-cell RNA sequencing data from metastatic melanoma specimens (GSE72056) (29) were analysed using the Seurat R package v5. Endothelial cells were identified according to their original cell-type annotations provided by Tirosh *et al.* and were used for downstream analysis. Endothelial cells were stratified according to detectable AMBRA1 expression, with cells exhibiting no detectable AMBRA1 expression classified as AMBRA1-negative (*n* = 32) and all remaining cells classified as AMBRA1-positive (*n* = 33). Module scores were calculated using Seurat `AddModuleScore()`. Three endothelial programmes were assessed: an IFN/antigen-presentation module (IFNAR1, IRF1, CIITA, HLA-A, HLA-B, HLA-C, B2M, TAP1 and TAP2), a leukocyte-recruitment module (VCAM1, ICAM1, SELPLG, CCL5, CCL22, TNFSF14 and TNFSF15), and an endothelial-identity module (JAM2, EGFL7, EMCN, FLT1, TEK, TIE1, ROBO4, CDH2, CXADR and KDR). Module scores were compared between AMBRA1-positive and AMBRA1-negative endothelial cell populations using two-sided Wilcoxon rank-sum tests. *P* values were adjusted (*padj*) for multiple testing using the Benjamini-Hochberg method. log_2_FC values were calculated by comparing mean expression between groups. Gene-wise Spearman analysis was performed using continuous AMBRA1 expression across endothelial cells and was used as the input for pre-ranked Hallmark GSEA. Hallmark pathways with a nominal *P* value < 0.05 and FDR < 0.25 were considered significant. Correlations between continuous AMBRA1 expression and IFN/antigen presentation, leukocyte recruitment and endothelial identity module scores were assessed using Spearman rank correlation analysis. For visualisation, and for direct comparison with the experimental siRNA-mediated AMBRA1 knockdown dataset, GSEA pathway rankings were inverted such that positive NES (NES > 0) represented pathways associated with low AMBRA1, and negative NES (NES < 0) represented pathways associated with high AMBRA1 expression.

### Western blotting

Cells were lysed with RIPA buffer supplemented with protease and phosphatase inhibitors (Protease Inhibitor Cocktail, 5 mM sodium fluoride, 0.5 mM sodium orthovanadate, 1 mM sodium molybdate and 0.5 mM phenylmethylsulfonyl fluoride; Sigma-Aldrich). Total protein extracts were separated using the 4-10% Mini-PROTEAN® TGX™ system (Bio-Rad) or hand-cast polyacrylamide gels (BioRad) and electroblotted onto nitrocellulose (BioRad or Whatman Amersham) or PVDF (Millipore) membranes, prior to incubation with primary antibodies in 5% non-fat dry milk in PBS (ThermoFisher Scientific) plus 0.1% Tween-20 (Sigma-Aldrich) overnight at 4°C (Supplementary Table 3). Detection was achieved using horseradish peroxidase–conjugated secondary antibodies (Cell Signalling Biotechnology, Jackson ImmunoResearch Laboratories; Supplementary Table 3) and enhanced chemiluminescence (ECL, Immobilon Western HRP substrate, Millipore; or Clarity™ Western ECL substrate, Bio-Rad). Signals were acquired using a ChemiDoc imaging system.

### Immunofluorescence

Cells were seeded onto sterile 20 mm x 20 mm glass cover slips (ThermoFisher) prior to treatment and fixation in 4% paraformaldehyde for 20 minutes at room temperature. Coverslips were treated with 4% bovine serum albumin (BSA) for 30 minutes at room temperature prior to incubation with anti-VE-cadherin overnight at 4 °C in 4% BSA, followed by anti-rabbit AlexaFluor® 546 for 1 hour at room temperature (Supplementary Table 3). Coverslips were mounted with DAPI-containing mounting media (Abcam) and imaged on a Leica DM6 B widefield microscope using SPOT Imaging software.

### Tubule formation assay

12-well flat bottom culture plates were coated with Matrigel® Matrix (Corning) as per the manufacturer’s instructions. Cells were seeded onto the solidified matrix at a density of 1.8 x10^5^ cells. Phase contrast images were taken using the EVOS™ XL Core inverted microscope (Thermo Fisher Scientific) at multiple timepoints post seeding, and analysed using the Angiogenesis Analyzer for Image J.

### Spheroid based adhesion assay

HUVEC cells were dissociated using 1X Versene (Gibco) to preserve cell-surface adhesion molecules, pelleted by gentle centrifugation and resuspended as a single-cell suspension before seeding into ultra-low attachment U-bottom 96-well plates (Costar) in 100 µL of culture medium. Cells were allowed to self-aggregate under gravity without centrifugation during spheroid formation and were imaged in real time using the IncuCyte™ live-cell imaging system (Sartorius) at hourly intervals up to 12 hrs. Object count (per phase contrast image) and aggregation dynamics were assessed using the Sartorius IncuCyte™ live-cell imaging software.

### Statistics

Unless stated otherwise, all experimental data are representative of the mean ±SEM of three independent biological replicates. Technical replicates within an experiment were averaged, apart from the tubule formation assay. Data were analysed by one sample *t-*test with Bonferroni correction for multiple comparisons, one-way or two-way ANOVA with appropriate post-hoc test for multiple comparisons, or by Mann-Whitney test for non-normally distributed data (GraphPad Prism v10 or R). Fold change (FC) data were log transformed prior to statistical analysis.

### Data availability

RNA sequencing (RNA-seq) data generated in this study are available in Gene Expression Omnibus (GEO) under accession number GSE342583. Associated processed data files are provided as supplementary data. All other data are available upon request from the corresponding author.

## Results

### Paracrine TGFβ signalling causes a reduction in endothelial AMBRA1 levels

We have previously demonstrated that loss of AMBRA1 and loricrin in the epidermis overlying non-ulcerated AJCC stage I/II melanomas are associated with risk of metastasis and that TGFβ2 signalling drives epidermal AMBRA1 loss (25,26). As endothelial cells contribute to the pro-metastatic microenvironment by supporting tumour growth and dissemination (10), we investigated whether TGFβ superfamily ligands also regulate endothelial AMBRA1. Analysis of the Cancer Genome Atlas Skin Cutaneous Melanoma (TCGA-SKCM) dataset (27) showed *TGFB1* and *TGFB3* expression were significantly higher in metastatic compared to primary melanomas (metastatic melanomas [*n* = 386], primary tumours [*n* = 104], Fig. 1A). To determine if tumour-derived factors regulate endothelial AMBRA1 *in vitro*, HMEC-1 cells were cultured in conditioned media derived from melanoma cell lines CHL-1 and A375, resulting in a significant reduction in AMBRA1 protein abundance at 72 hrs, but not 24 hrs, while AMBRA1 mRNA levels remained unchanged (Fig. 1B). To assess whether TGFβ ligands directly mediated this effect, primary and immortalised endothelial cells were stimulated with individual TGFβ1, TGFβ2 or TGFβ3 isoforms for 24 or 72 hrs, which induced a dose-dependent reduction in AMBRA1 protein levels in HUVEC, HDBEC and HMEC-1 cells at 72 hrs, but not at 24 hrs, without affecting AMBRA1 transcript levels (Fig. 1C; Supplementary Fig. S1,S2;). Combined treatment with pooled TGFβ1/2/3 ligands also significantly reduced AMBRA1 protein levels in HUVEC and HDBEC endothelial cells (Fig. 1D). Given endothelial AMBRA1 loss occurred in response to melanoma-derived factors and specifically sustained TGFβ exposure, we next sought to define the global transcriptional and cellular consequences of endothelial AMBRA1 loss.

**Figure 1.**
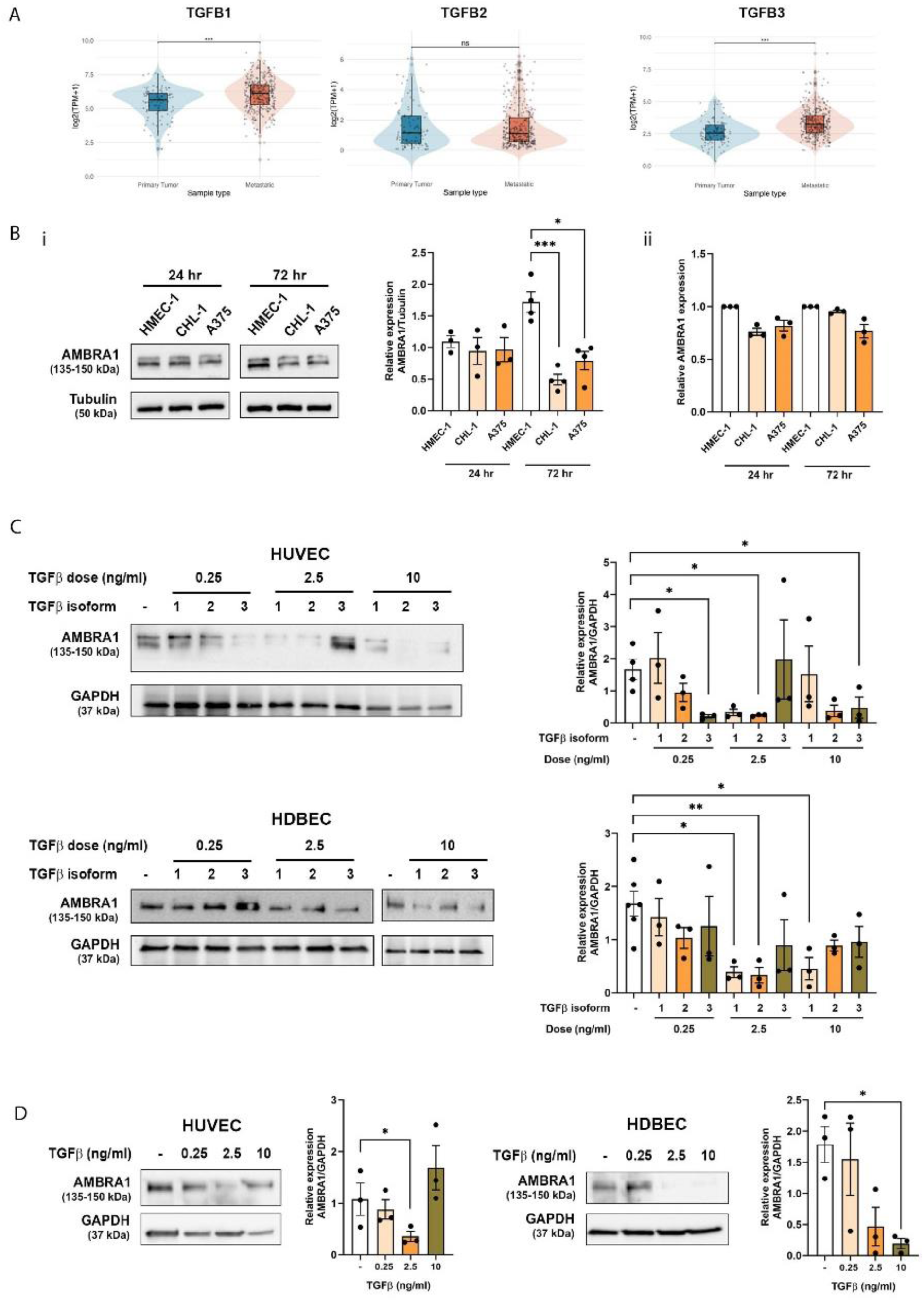
TGFβ-mediated signalling drives loss of endothelial AMBRA1 in a concentration and context dependent manner. (A) TGFB1, TGFB2 and TGFB3 transcript expression in metastatic melanomas (n = 368) compared with primary tumours (n = 104) from The Cancer Genome Atlas Skin Cutaneous Melanoma (TCGA-SKCM) dataset. (B) HMEC-1 cells were exposed to conditioned medium from HMEC-1, CHL-1 or A375 cells for 24 or 72 hrs. (i) AMBRA1 and tubulin protein abundance were quantified by densitometry, normalised to tubulin and presented relative to the mean protein/tubulin value for each experiment and time point (mean ± SEM, n ≥ 3; one-way ANOVA with Sidak’s multiple comparison test on logFC data; * *P* < 0.05, *** *P* < 0.001). (ii) mRNA expression levels were normalised to RPL34 and presented relative to siNT (mean ± SEM, n = 3). (C,D) Western blots of AMBRA1 and GAPDH protein from HUVEC and HDBEC cells in (C) the absence or presence of treatment with TGFβ 1, 2 or 3 (0.25-10 ng/ml) for 72 hrs, or (D) with pooled TGFβ isoforms (0.25-10 ng/ml) for 72 hrs. Protein levels were quantified by densitometry, normalised to GAPDH and presented relative to the mean protein/GAPDH value for each experiment (mean ± SEM, *n* = 3; one-way ANOVA with Dunnett’s multiple comparison test on logFC data to compare treatment to control; * *P* < 0.05, ** *P* < 0.01).

### Loss of AMBRA1 activates cell-cycle programmes and suppresses endothelial adhesion, inflammatory signalling, ECM engagement, and angiogenic identity

To define the transcriptional consequences of endothelial AMBRA1 loss, RNA-sequencing was performed in HUVEC cells following siRNA-mediated depletion. AMBRA1 knockdown was confirmed prior to sequencing (Supplementary Fig. S3A). AMBRA1 depletion resulted in 11,739 DEGs, of which 488 demonstrated statistically significant differential expression (*padj* < 0.05) and 39 showed a statistically significant 1.5-fold change (log_2_ fold change [log_2_FC] ± 0.585, *padj* < 0.05), including AMBRA1 itself (Supplementary Fig. S3B-C; Supplementary Table 5).

Gene set enrichment analysis demonstrated activation of proliferative programmes, including MYC Targets, G2M Checkpoint, E2F Targets and MTORC1 Signalling, together with enrichment of GO terms related to mitosis and chromosome segregation (Fig. 2A,B; Supplementary Fig. S4A; Supplementary Tables 6-8). Conversely, AMBRA1 loss was associated with suppression of pathways governing adhesion, extracellular matrix organisation and angiogenic function, including Focal Adhesion, Cell Adhesion Molecules (CAMs) and ECM-Receptor Interaction, together with suppression of GO terms associated with endothelial structural integrity, Cell–Cell Junction and Blood Vessel Morphogenesis (Fig. 2B,C; Supplementary Fig. S4B; Supplementary Tables 7-8). Consistent with these pathway-level changes, several endothelial-associated genes were significantly reduced following AMBRA1 depletion, including adhesion molecules *CDH2* (N-cadherin) and *CXADR*, ECM anchoring genes *LAMC2*, *AGRIN* and *COLGALT1,* and the angiogenic regulator *NOTCH4* (Fig. 2D).

**Figure 2.**
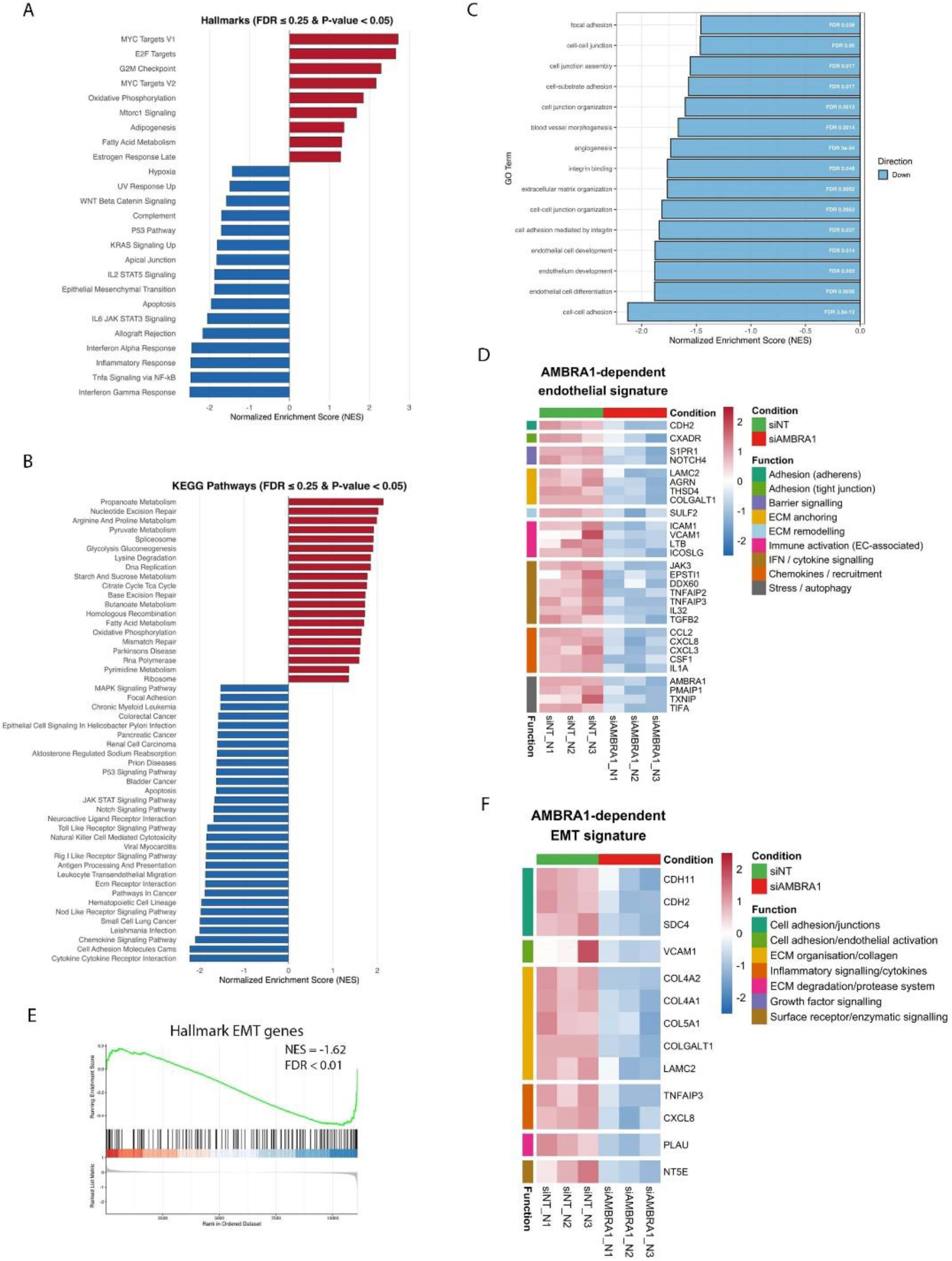
AMBRA1 loss suppresses cell-cell adhesion, immune-response and endothelial programmes in vitro, correlated with endothelial cell dysfunction. (A-F) RNA-sequencing was performed on HUVEC cells following siRNA-mediated AMBRA1 depletion or non-targeting (siNT) control for 48 hrs. (A,B) DEGs were analysed against Hallmark (A) and KEGG (B) gene sets. Graphs shown the top 15 (or less) positively and negatively enriched pathways with a P value < 0.05 and an FDR < 25%. (C) Gene Ontology enrichment analysis of DEGs related to adhesion, cytoskeletal organisation and angiogenesis (*padj* < 0.05). (D) Heatmap of significantly differentially expressed genes involved in endothelial adhesion, extracellular matrix (ECM) remodelling, barrier regulation, and immune-associated signalling (*padj* < 0.05). (E) GSEA using the Hallmark epithelial-to-mesenchymal transition (EMT) gene signature, with enrichment significance assessed using P-value < 0.05. (F) Heatmap of significantly differentially expressed leading-edge genes contributing to EMT gene set enrichment (*padj* < 0.05).

AMBRA1 loss was also associated with broad suppression of inflammatory and immune-response programmes. Hallmark and KEGG analyses demonstrated significant negative enrichment of IFNγ Response, IFNα Response, TNFα Signalling via NFκB, Inflammatory Response, Cytokine-Cytokine Receptor Interaction, Chemokine Signalling, and Leukocyte Trans-Endothelial Migration (Fig. 2A,B; Supplementary Tables 6-7). Several mediators of endothelial inflammatory activation and leukocyte recruitment, including *ICAM1*, *VCAM1*, *CXCL8*, *CXCL3*, *CCL2*, *IL1A* and *TNFAIP3,* were significantly downregulated (Fig. 2D), suggesting endothelial AMBRA1 loss may impair immune-cell trafficking.

Given the prominent transcriptional changes in adhesion, ECM organisation and endothelial identity programmes, we next assessed Hallmark Epithelial–Mesenchymal Transition (EMT). EMT was significantly negatively enriched following AMBRA1 depletion (NES = -1.62, FDR = 0.0049) (Fig. 2A,E; Supplementary Table 6). Analysis of significantly differentially expressed (*padj* < 0.05) leading edge EMT-associated genes showed reduced expression of junctional and ECM-associated genes, including *CDH2/11*, *SDC4*, *COL4A1/2*, *COL5A1* and *LAMC2*, alongside reduced expression of inflammatory genes such as *TNFAIP3*, *PLAU* and *CXCL8* (Fig. 2F). Collectively, these data suggest endothelial AMBRA1 loss drives transcriptional changes governing proliferation, adhesion and vascular organisation, promoting further investigation into endothelial identity and EndoMT-associated pathways.

### Endothelial AMBRA1-associated transcriptional programmes are recapitulated in human melanoma vasculature

To determine whether the transcriptional phenotype associated with endothelial AMBRA1 loss *in vitro* was also evident in human melanoma vasculature, we analysed endothelial cells from a publicly available melanoma single-cell RNA sequencing dataset generated by Tirosh *et al.* (GSE72056) (29), comprising 4,645 cells isolated from 19 human metastatic melanoma specimens. Endothelial cells (*n* = 65) were identified from the original annotations and stratified by detectable AMBRA1 expression, with 33 AMBRA1-positive and 32 AMBRA1-negative cells.

Module scores for IFN/antigen presentation, leukocyte recruitment and endothelial identity programmes were significantly increased in AMBRA1-expressing endothelial cells across all three programmes compared with AMBRA1-negative cells (Fig. 3A), suggesting reduced endothelial AMBRA1 expression is associated with loss of immune-supportive and endothelial-stabilising vascular functions. AMBRA1-positive cells exhibited increased expression of endothelial-associated genes, including *FLT1*, *EGFL7*, *EMCN*, *JAM2*, and *CDH2*, together with genes involved in inflammatory activation and leukocyte recruitment, including *VCAM1*, *ICAM1*, *SELPLG*, *IFNAR1*, *CIITA*, *CCL5*, *CCL22*, *TNFSF14*, and *TNFSF15* (Fig. 3Bi; Supplementary Table 9).

**Figure 3.**
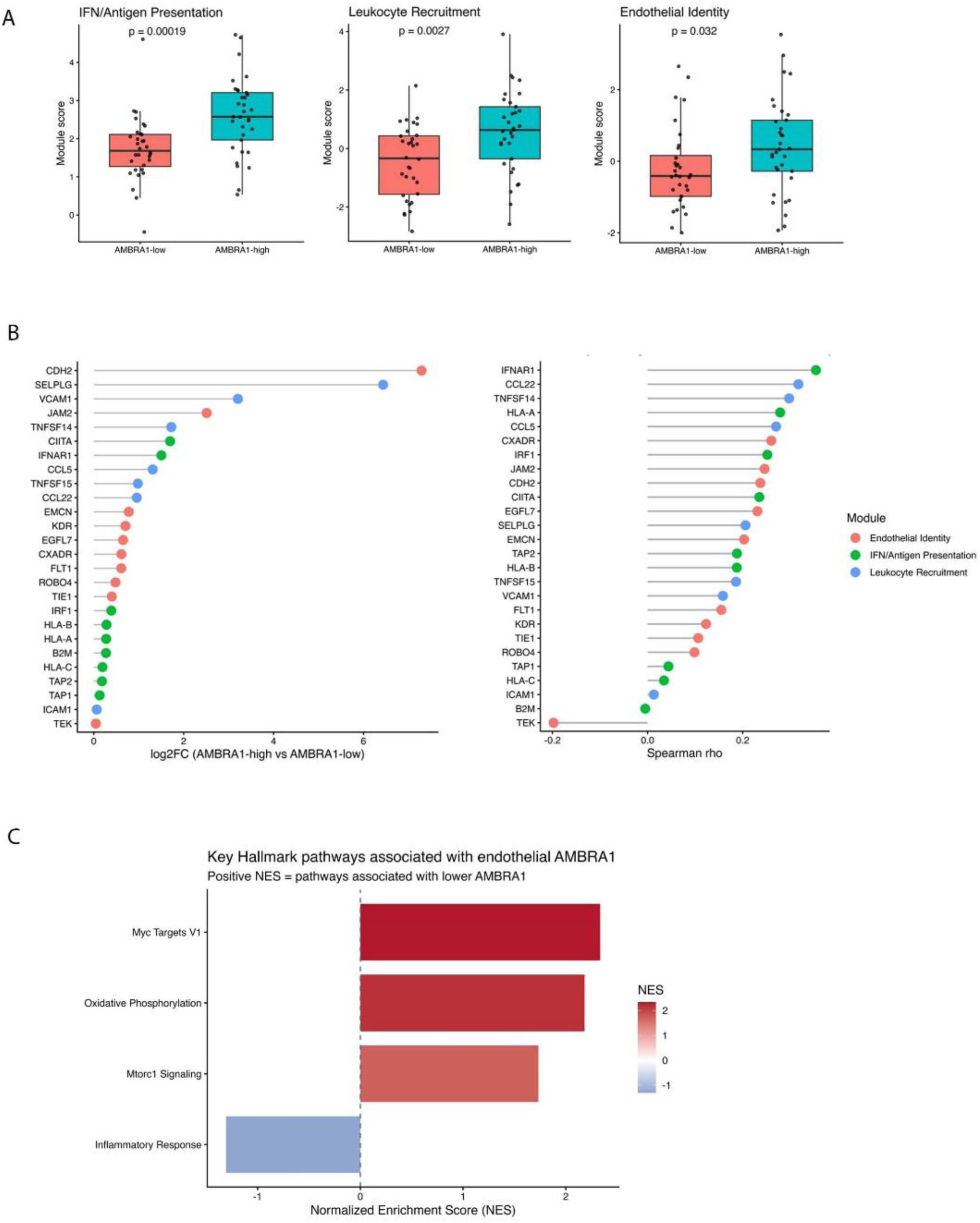
Endothelial AMBRA1-associated transcriptional programmes in human melanoma vasculature. (A) Module scores for IFN/antigen presentation genes (IFNAR1, IRF1, CIITA, HLA-A, HLA-B, HLA-C, B2M, TAP1 and TAP2), leukocyte recruitment genes (VCAM1, ICAM1, SELPLG, CCL5, CCL22, TNFSF14 and TNFSF15), and endothelial identity genes (JAM2, EGFL7, EMCN, FLT1, TEK, TIE1, ROBO4, CDH2, CXADR and KDR) in AMBRA1-positive and AMBRA1-negative endothelial cells (two-sided Wilcoxon rank-sum test). (B) log_2_FC in expression of selected IFN/antigen presentation, leukocyte recruitment and endothelial identity genes in AMBRA1-negative and AMBRA1-positive endothelial cells. Average expression was calculated separately across all cells within each group prior to log_2_FC calculation, or (right) Spearman correlation coefficients for genes included in the IFN/antigen presentation, leukocyte recruitment and endothelial identity programmes across all endothelial cells using continuous AMBRA1 expression values. Positive values indicate genes whose expression increases with AMBRA1 expression. (C) Hallmark GSEA performed on genes ranked according to their Spearman correlation coefficient with AMBRA1 expression. Rankings were inverted such that positive NES (NES > 0) represent pathways associated with low AMBRA1 expression, and negative NES (NES < 0) represent pathways associated with high AMBRA1 expression. Pathways were filtered using the conventional GSEA significance thresholds of nominal P < 0.05 and FDR < 0.25.

To account for the binary stratification of endothelial cells, gene-wise Spearman correlation analyses were additionally performed using continuous AMBRA1 expression values across all endothelial cells (Supplementary Table 10). Consistent with the module score analysis, IFN/antigen presentation, endothelial identity and leukocyte recruitment programme scores were significantly positively correlated with continuous AMBRA1 expression (Supplementary Table 11). Similar associations were observed at the individual gene level (Fig. 3Bii), supporting a relationship between endothelial AMBRA1 expression and immune-supportive programmes. To determine whether these gene-level associations translated into broader transcriptional programmes, genes were ranked according to their Spearman correlation coefficient with AMBRA1 expression and subjected to Hallmark GSEA. Consistent with the transcriptional consequences of siRNA-mediated AMBRA1 depletion, low endothelial AMBRA1 expression was associated with enrichment of proliferative programmes, including MYC Targets and MTORC1 signalling, together with suppression of inflammatory response and KRAS signalling pathways in both datasets (Fig. 3C, Supplementary Table 12). Together, these findings indicate that endothelial transcriptional states associated with siRNA-mediated AMBRA1 loss are recapitulated within human melanoma vasculature and are characterised by suppression of endothelial identity, inflammatory signalling and leukocyte-recruitment programmes.

### AMBRA1 loss disrupts endothelial identity without inducing canonical EndoMT

Given suppression of adhesion- and ECM-associated gene programmes following endothelial AMBRA1 loss (Fig. 2D-E), we next evaluated whether AMBRA1 loss drives bona fide EndoMT. As no curated EndoMT signature specific to oncology and skin vasculature was available, a bespoke EndoMT gene signature was curated from published studies, incorporating endothelial identity markers, EndoMT inducers, transcriptional regulators and mesenchymal markers (Supplementary Table 4). Benchmarking against a published EndoMT dataset generated from TGFβ2-treated HUVECs (GSE118446) (30), demonstrated significant enrichment (NES = 1.64, FDR = 0.00135), confirming sensitivity to detect EndoMT-associated transcriptional changes (Supplementary Fig. S5A, B).

GSEA revealed negative enrichment of the EndoMT signature following AMBRA1 depletion in HUVECs (NES = -1.89, FDR < 0.001) (Fig. 4A), indicating AMBRA1 loss does not induce canonical EndoMT. Instead, AMBRA1 depletion was associated with broad suppression of endothelial junction identity genes, including *CLDN5* (Claudin-5), *CDH5*, *F11R*, *DCBLD2*, *TEK* and *TIE1*, while only two endothelial identity-associated markers (*NOS3* and *LYVE1*) were increased. Additionally, several endothelial-associated inflammatory and immune-supportive genes, including *VCAM1* and *ICAM1*, were also downregulated, whereas genes associated with metabolic and stress rewiring, including *FABP5*, *DHCR24* and *HMGCS1*, were upregulated. Consistent with the negative enrichment of the EndoMT signature, upstream EndoMT-inducers (*ACVRL1*, *IL1A*, *TGFB2*, *TGFBR2*) and transcriptional drivers (*NFKB2*, *RELA*, *RELB*) were reduced. Importantly, mesenchymal effector genes were not induced, with *CDH2* (N-cadherin) and *CNN2* significantly decreased (Fig. 4B). Consistent with these transcriptional changes, endothelial AMBRA1 depletion reduced expression of tight-junction protein Claudin-5 (CLDN5) at both the protein and transcript levels in HMEC-1 cells (Fig. 4C), reduced N-cadherin protein abundance (Fig. 4D) and decreased cell surface VE-cadherin expression in HUVECs (Fig. 4E). Collectively, these data suggest TGFβ-mediated AMBRA1 loss compromises endothelial junctional integrity and vessel stabilisation without inducing full EndoMT.

**Figure 4.**
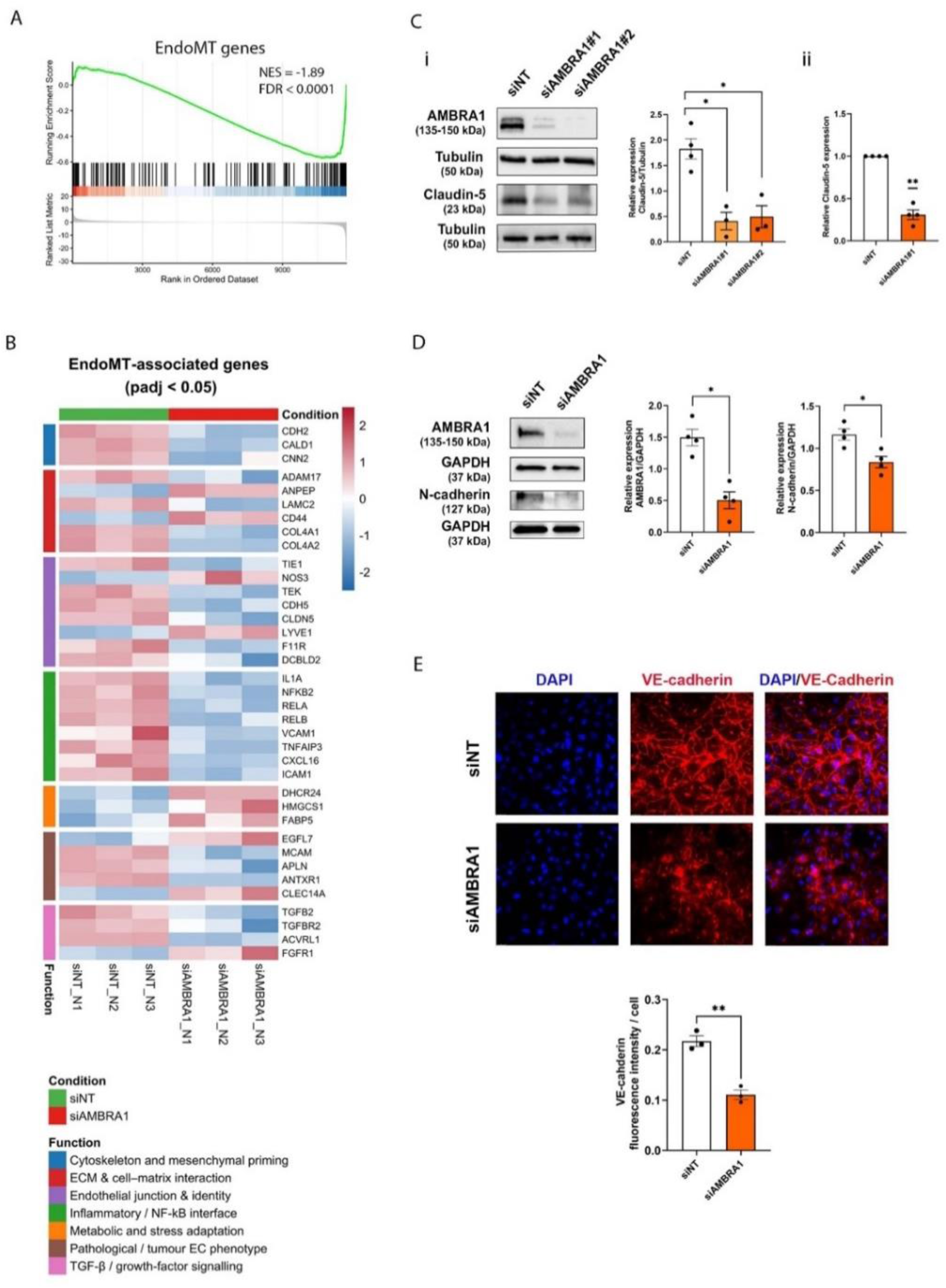
AMBRA1 loss disrupts endothelial identity without canonical EndoMT. (A) GSEA of DEGs from AMBRA1-depleted HUVEC cells compared with siNT controls using a bespoke EndoMT gene signature (B) Heatmap of significantly differentially expressed genes contributing to EMT gene set enrichment (*padj* < 0.05). (C) HMEC-1 or (D-E) HUVEC cells were transfected with control (siNT) or AMBRA1 siRNA for 48 hrs. (C) (i) AMBRA1, Claudin-5 and tubulin protein levels were quantified by densitometry, normalised to tubulin and presented relative to the mean Claudin-5/tubulin value for each experiment (mean ± SEM, *n* = 3; one-way ANOVA with Dunnett’s multiple comparison test on log FC data; * *P* < 0.05). (ii) mRNA expression levels were normalised to RPL34 and presented relative to siNT (ean ± SEM, *n* = 3; one-sample *t*-test on logFC data; ** *P* < 0.01). (D) AMBRA1, N-cadherin and GAPDH protein levels were quantified by densitometry, normalised to GAPDH and presented relative to the mean N-cadherin/GAPDH value for each experiment (mean ± SEM, *n* = 4) (*t*-test on logFC data; * *P* < 0.05). (E) Immunofluorescence analysis of VE-cadherin. Data are mean VE-cadherin fluorescence intensity per image, normalised to cell number (mean ± SEM of 3 technical triplicates across *n* = 3; *t*-test, ** *P* < 0.01).

### AMBRA1 loss promotes early tubulogenesis and endothelial cell proliferation

To assess the functional consequences of endothelial AMBRA1 loss, we depleted AMBRA1 in HMEC-1 and HUVEC cells and assessed their ability to form capillary-like structures in a Matrigel matrix. AMBRA1 knockdown accelerated early tubulogenesis, with a significant increase in the number of master junctions, segments and meshes, and mean mesh size at 4 hrs in HUVECs, and from 90 minutes through to 7 hrs in HMEC-1 cells (Fig. 5A,B; Supplementary Fig. 6A,B). siRNA-mediated depletion of AMBRA1 in HUVECs also resulted in a significant increase in cell proliferation (Fig. 5C,D), consistent with AMBRA1 loss promoting activation of transcriptional programmes associated with cell cycle progression. Furthermore, AMBRA1-depleted HUVECs failed to form compact spheroids over 12 hrs, with a significantly higher number of discrete objects following AMBRA1 knockdown compared to control cells (Fig. 5E),, consistent with reduced adhesion- and junction-associated pathways and proteins. Together, these data suggest AMBRA1 loss increases endothelial cell proliferation and accelerates early tubulogenesis, consistent with transcriptomic changes indicative of a hyperproliferative endothelial state, whilst simultaneously compromising endothelial cell-cell adhesion and cellular assembly.

**Figure 5.**
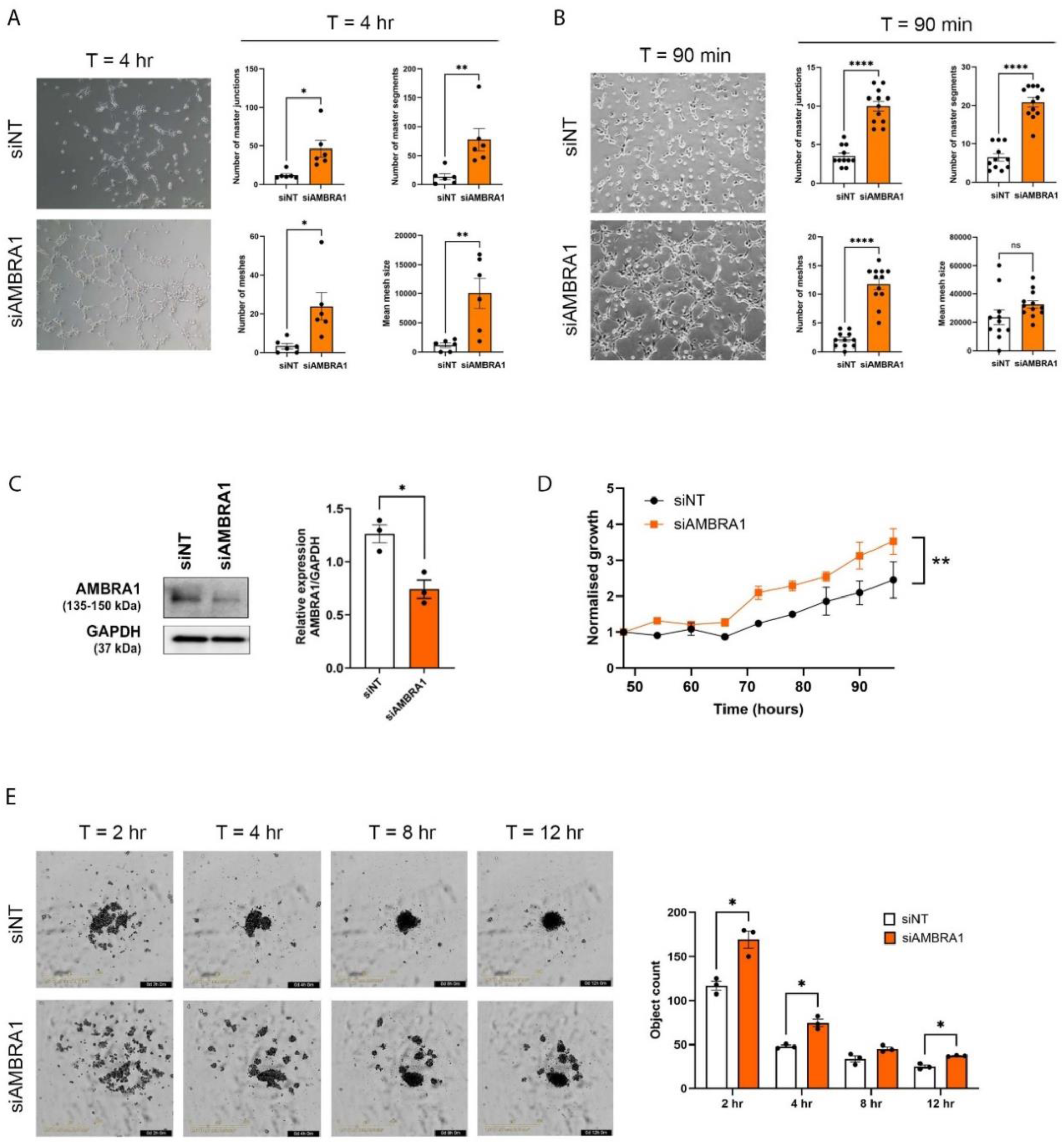
AMBRA1 knockdown enhances endothelial cell vascular network formation. (A,B) HUVEC cells transfected with control (siNT) or AMBRA1 siRNA (A) or HMEC-1 cells were transfected with control (siNT) or AMBRA1 #1 siRNA (B) for 48 hrs were seeded onto Matrigel and tubule formation monitored over time. Images were taken using a 10X objective, and the number of master junctions, master segments and meshes, and mean mesh size, were quantified at 4 hr (A) or 90 min (B) (mean ± SEM, each data point is a technical replicate from a minimum of two independent experiments; *t*-test, * *P* < 0.05). (C,D) Western blots of AMBRA1 and GAPDH protein from HUVEC cells transfected with control (siNT) or AMBRA1 siRNA for 48 hrs. Protein levels were quantified by densitometry, normalised to GAPDH and presented relative to the mean AMBRA1/GAPDH value for each experiment (mean ± SEM, *n* = 3) (*t*-test on logFC data; * *P* < 0.05). Cell proliferation was monitored from 48 to 96 hrs and normalised to growth at 48 hr (repeated measures two-way ANOVA on log FC data; ** *P* < 0.01). (E) HUVEC cells treated with control (siNT) or AMBRA1 siRNA for 48 hrs were seeded into ultra-low attachment U-bottom plates and were imaged in real time using the IncuCyte™ live-cell imaging system (Sartorius) at 2, 4, 6 and 12 hrs (mean ± SEM of 3 technical triplicates across *n* = 3; *t*-test, * adj. *P* < 0.05) (scale bar = 800 mm).

### EndoMT regulators Snail and Slug are elevated by the tumour secretome and endothelial AMBRA1 loss

Given that AMBRA1 loss altered endothelial identity and junctional integrity without inducing complete canonical EndoMT, we investigated whether key EndoMT-associated transcriptional regulators Slug and Snail, downstream effectors of TGFβ signalling (31), were altered following endothelial AMBRA1 loss. In HUVEC cells, AMBRA1 knockdown produced a modest but statistically significant increase in Snail protein (Fig. 6A), whereas Slug protein was not detectable. In contrast, AMBRA1 depletion in HMEC-1 cells significantly increased both Snail and Slug protein abundance (Fig. 6A), without a corresponding increase in mRNA expression (Fig. 6B). Accumulation of Slug following treatment with MG132, but not bafilomycin A1, suggested proteasome-mediated regulation (Fig. 6C). We next assessed if melanoma-derived factors influence endothelial Snail protein expression. Melanoma-conditioned medium from CHL-1 and A375 cells induced a significant increase in Snail transcript at 24 hrs, but not at 72 hrs, in HMEC-1 cells, whereas Snail protein remained elevated at both timepoints (Fig. 6D). Finally, given AMBRA1 can function as a substrate receptor within the DDB1-CUL4 E3 ubiquitin ligase complex, we assessed whether this pathway contributes to Snail regulation. siRNA-mediated DDB1 knockdown did not alter Snail protein abundance in HMEC-1 cells (Fig. 6E). Collectively, these data show that AMBRA1 loss stabilises the EndoMT master regulators Slug and Snail.

**Figure 6.**
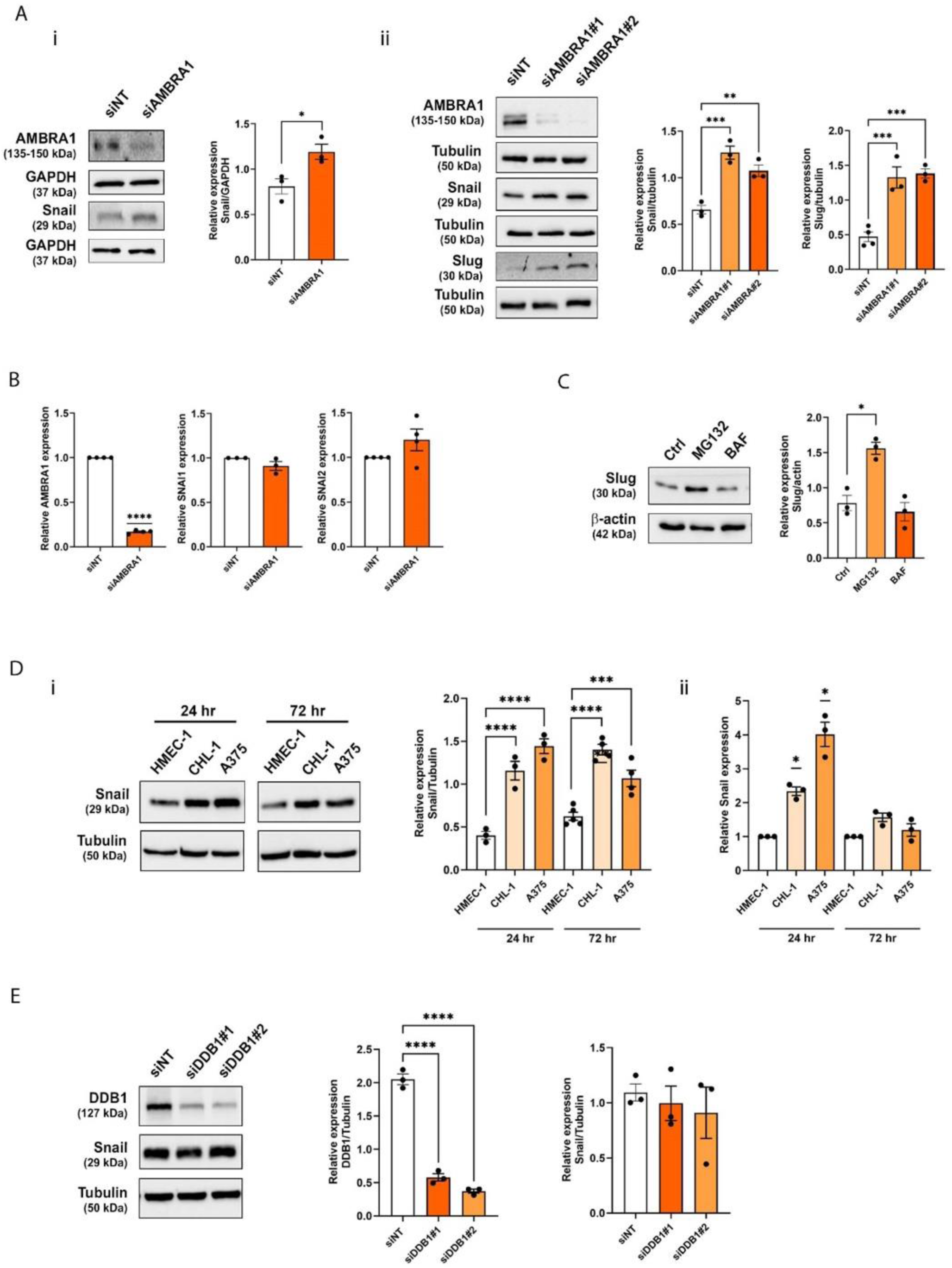
AMBRA1 regulates master EMT regulators Snail and Slug. (A) Western blots of AMBRA, Snail and GAPDH from HUVEC cells (i) and AMBRA1, Snail, Slug and tubulin from HMEC-1 cells (ii) transfected with control (siNT) or AMBRA1 siRNA for 48 hrs. (B) qPCR mRNA expression analysis of AMBRA1, SNAI1, SNAI2 and RPL34 from HMEC-1 cells transfected with control (siNT) or AMBRA1 siRNA #1 for 48 hrs. (C) Western blots of Snail and β-actin from HMEC-1 cells treated with MG132 (5 µM) or Bafilomycin A1 (BAF; 5 nM) for 4 hrs. (D) Western blots (i) and mRNA analysis (ii) of Snail and tubulin or RPL34 in HMEC-1 cells exposed to conditioned medium obtained from HMEC-1, CHL-1 or A375 cells for 24 or 72 hrs. (E) Western blot of DDB1, Snail and tubulin from HMEC-1 cells transfected with control (siNT) or DDB1 siRNA for 48 hrs. Protein levels were quantified by densitometry, normalised to the loading control and presented relative to the mean protein/loading control value for each experiment (mean ± SEM, n ≥ 3) (*t* test or one-way ANOVA with Dunnett’s or Sidak’s multiple comparison test on log FC data to compare treatment to control; * P < 0.05, ** P < 0.01, *** P < 0.001, **** P < 0.0001), and mRNA expression levels were normalised to RPL34 and presented relative to siNT (mean ± SEM, n = 3) (one-sample *t*-test on logFC data; * adj. *P* < 0.05).

### Loss of endothelial AMBRA1 is associated with increased risk of metastasis in early-stage melanoma

To determine the clinical relevance of endothelial AMBRA1 loss in early-stage melanoma, we examined AMBRA1 immunohistochemical (IHC) staining within the tumour-associated vasculature in a cohort of 121 AJCC stage I/II primary melanomas previously characterised as AMBLor at-risk based on loss of AMBRA1 and loricrin expression in the peritumoural epidermis. Endothelial AMBRA1 expression was assessed in blood and lymphatic vessels within the tumour body (intratumoural vessels) and within 500 µm of the tumour boundary (peritumoural vessels), relative to the average vascular AMBRA1 expression in vessels across the individual tissue section (including the dermis and margin; Fig. 7A).

**Figure 7.**
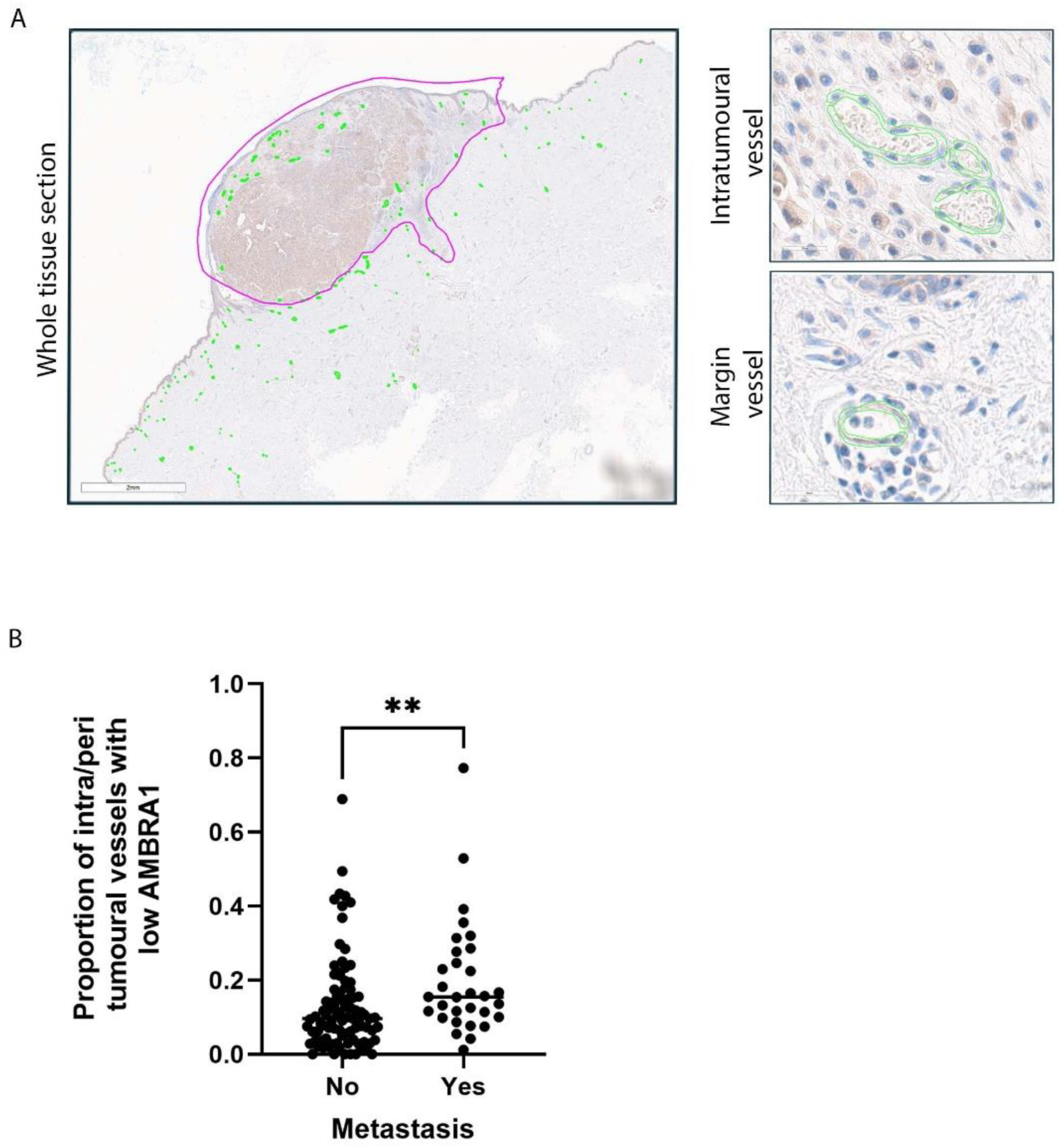
Decreased endothelial AMBRA1 is associated with metastasis of early-stage melanoma. (A) Representative section of a non-ulcerated AJCC stage I/II melanoma stained by immunohistochemistry (IHC) for AMBRA1, annotated for the tumour boundary (purple) and blood/lymphatic vessels (green), with a magnified view of annotated intratumoural and marginal vessels (scale bars 2 mm, 30 mm). (B) The proportion of intra- and peri-tumoural vessels with low AMBRA1 expression (< 0.45 x average AMBRA1 score) was determined in a cohort of non-ulcerated AJCC stage I/II melanomas (*n* = 121), with tumours grouped according to subsequent metastatic outcome. Horizontal line represents the median (Mann-Whitney test; ** *P* < 0.01).

Patients with primary melanoma who later experienced tumour recurrence (in-transit [*n* = 3], nodal [*n* = 12] or distant [*n* = 16] metastases) exhibited a significant increase in the proportion of intratumoural and peritumoural vessels with low levels of AMBRA1, compared to patients whose tumours did not recur (*P* < 0.01; Fig. 7B), suggesting endothelial AMBRA1 loss is associated with metastatic spread.

## Discussion

Despite advances in targeted and immune therapies, melanoma remains the leading cause of skin cancer-related mortality (1). Current AJCC staging criteria fails to capture early-stage AJCC stage I/II melanoma patients at greatest risk of recurrence and metastatic progression, highlighting the need for biologically-informed prognostic biomarkers capable of refining risk stratification beyond conventional clinicopathological features (32). We previously showed that loss of AMBRA1 and loricrin in the epidermis overlying non-ulcerated AJCC stage I/II melanomas identifies a subset of patients at increased risk of recurrence (25). Here, we extend the tumour-suppressive role of AMBRA1 beyond the epidermis and characterise endothelial AMBRA1 as a regulator of vascular dysfunction and remodelling, supporting the concept that vascular alterations contribute to early disease progression (33,34).

Given we have previously shown TGFβ2 drives AMBRA1 loss in the overlying epidermis of AJCC stage I/II melanomas (26), we investigated whether TGFβ signalling drives a similar mechanism within the tumour-associated endothelium. We show *TGFB1* and *TGFB3* expression is elevated in metastatic versus primary melanomas, consistent with previous reports implicating TGFβ-signalling in melanoma progression (35,36). Furthermore, melanoma-conditioned media and prolonged TGFβ exposure reduced endothelial AMBRA1 protein abundance without altering transcript levels, suggesting chronic tumour-derived TGFβ may promote post-transcriptional endothelial AMBRA1 loss, consistent with the established ability of sustained TGFβ to modulate endothelial identity and protein stability (33,36).

Endothelial AMBRA1 loss in HUVECs revealed transcriptional reprogramming characterised by activation of proliferative programmes, consistent with the established role of AMBRA1 in restraining cell-cycle progression (11,15,17,37–39), and the coordinated suppression of endothelial adhesion, junctional organisation, and immune-supportive transcriptional programmes. Importantly, these observations were recapitulated in an independent human melanoma single-cell RNA sequencing dataset (29), in which low endothelial AMBRA1 expression was associated with reduced antigen presentation, leukocyte recruitment and endothelial identity programmes. Endothelial cells regulate vascular integrity and immune-cell trafficking through expression of adhesion molecules, chemokines and antigen-presentation machinery (12, 13). Suppression of these programmes may impair immune surveillance and promote immune exclusion. As endothelial dysfunction is increasingly recognised as a determinant of tumour progression and therapeutic responsiveness (16, 17), our findings suggest endothelial AMBRA1 loss may influence melanoma progression through vascular destabilisation and impaired immune-endothelial interactions. This phenotype is consistent with dysfunctional tumour vasculature, whereby endothelial proliferation becomes uncoupled from proper junctional organisation and vessel maturation, resulting in increased permeability, tumour cell intravasation and dissemination (8,33).

Although many of the adhesion- and ECM-associated genes downregulated in response to AMBRA1 loss are typically linked to cellular plasticity, further analysis showed this did not correspond with activation of classical EMT or canonical EndoMT programmes. Instead, AMBRA1 depletion resulted in selective suppression of endothelial identity markers, including *CDH5* (VE-cadherin), *CDH2* (N-cadherin), *CLDN5* (Claudin-5), *TIE* and *TEK*, without induction of mesenchymal effector genes, consistent with the emerging concept of partial or non-canonical endothelial plasticity, whereby endothelial cells lose aspects of cellular identity and junctional stability markers without fully acquiring mesenchymal fate (31,40–42). These states have been linked to increased permeability and barrier dysfunction (43), altered transendothelial trafficking (43,44) and tumour dissemination (44,45), whilst retaining sufficient endothelial characteristics to support vascularisation (46). Importantly, these transcriptomic changes were accompanied by downregulation of key junctional components Claudin-5, a critical determinant of endothelial barrier integrity, (33,47), and VE- and N-cadherins, which mediate endothelial-endothelial and endothelial-mural cell interactions, respectively (48–52). Loss or dysregulation of VE-cadherin disrupts junction stability and barrier function (48–50), while reduced endothelial N-cadherin is linked to impaired pericyte attachment and vessel maturation (51,52). Together, these findings suggest endothelial AMBRA1 loss impairs vascular stability through disruption of endothelial junctions and mural support rather than induction of full EndoMT.

Despite the absence of canonical EndoMT, AMBRA1 loss was associated with post-transcriptional increases in the EndoMT-associated transcription factors Slug and Snail. Consistent with previous reports demonstrating post-translational regulation of Slug and Snail (53,54), our data suggest proteasomal inhibition and tumour-derived factors can influence the stability of these transcription factors down stream of endothelial AMBRA1 loss. Although AMBRA1 can functions as a substrate receptor for the Cullin4-DDB1-RBX1 E3 ligase complex (13), depletion of DDB1 did not result in Snail accumulation; however, this does not exclude a role for AMBRA1 in the regulation of Slug/Snail stabilisation through interactions with components of other E3 ligase complexes (13). Although classically associated with EMT and EndoMT (31,40), partial Slug/Snail activation can alter vascular adhesion and barrier organisation without full mesenchymal transition (40,55). Our findings suggest endothelial AMBRA1 loss promotes endothelial plasticity characterised by junctional destabilisation and altered vascular function without progression to a complete mesenchymal fate.

Functionally, endothelial AMBRA1 loss promoted accelerated early tubulogenesis, increased proliferation and impaired cell-cell adhesion, consistent with an immature, disorganised vasculature phenotype. Given links between aberrant vascular remodelling, metastasis, immune exclusion and impaired therapeutic delivery (7–10,33,34), endothelial AMBRA1 loss may represent a previously unrecognised contributor to early-stage melanoma progression. Consistent with this, we demonstrate that AMBRA1 loss within tumour-associated blood and lymphatic endothelium surrounding AJCC stage I/II melanomas is significantly associated with increased metastatic risk. Whilst association does not establish causality, it is consistent with our experimental findings showing endothelial AMBRA1 loss promotes vascular dysfunction, impaired junctional integrity and suppression of immune-supportive programmes. These findings suggest vascular alterations may precede clinically detectable metastatic spread, supporting the concept that endothelial dysfunction is not merely a consequence of advanced disease but may also contribute to the establishment of a pro-metastatic environment.

Several limitations of this study should be acknowledged. First, the patient cohort was retrospective and intentionally enriched for recurrence events, (31/121 cases, ∼25%). While this proportion is higher than would be expected in an unselected early-stage melanoma cohort, it remains within the reported range of recurrence rates across stage I-II disease (10–20% for IA/IB; ∼30–59% for IIA/IIB/IIC) (6). Importantly, endothelial AMBRA1 assessment was performed blinded to recurrence outcome, reducing observer bias. Secondly, validation of the endothelial AMBRA1-associated transcriptional programmes utilised a transcriptomic data from metastatic melanoma samples. Therefore, the endothelial transcriptional programmes identified may reflect additional vascular remodelling associated with advanced disease. However, concordance between the single-cell dataset and transcriptional changes observed following experimental AMBRA1 loss supports the relevance to human melanoma vasculature. Finally, whilst our data support a link between tumour-derived signalling, endothelial AMBRA1 loss and vascular dysfunction, *in vivo* modelling will be necessary to fully define how endothelial AMBRA1 contributes to melanoma progression and metastatic dissemination.

In summary, this study identifies endothelial AMBRA1 as a key regulator of vascular integrity and plasticity in early-stage melanoma, linking tumour-derived TGFβ signalling to junctional destabilisation, vascular dysfunction, transcriptional reprogramming, and subsequent metastatic risk for patients with AJCC I/II melanomas. These findings further expand the role of AMBRA1 within the melanoma microenvironment and highlight the tumour vasculature as under-recognised component of early-stage melanoma progression, with potential implications for further refining risk stratification beyond conventional clinicopathological staging.

## Acknowledgments

For the purpose of open access, the author has applied a Creative Commons Attribution (CC BY) licence to any Author Accepted Manuscript version arising from this submission.

## Author contributions

LW conducted most of the experiments, together with RoS, and analysed and curated the data. MDR, KB, SB, LO and IC performed some experiments. LW, along with KB, performed all bioinformatics. PS contributed to validation of histopathology samples, and GR was involved in project administration. RaS, PEL and JA supervised the project. MP, GMF, PEL and JA devised the project, whilst KB, RaS, GMF, PEL and JA acquired funding for the work. LW, PEL and JA wrote the original draft, and LW, MDR, KB, MP, GMF, PEL and JA reviewed and edited the article.

