## Supplementary material for "Endothelial AMBRA1 loss contributes to vascular dysregulation facilitating metastatic progression in early-stage melanoma"

**Supplemental material**

Supplementary Figure 1

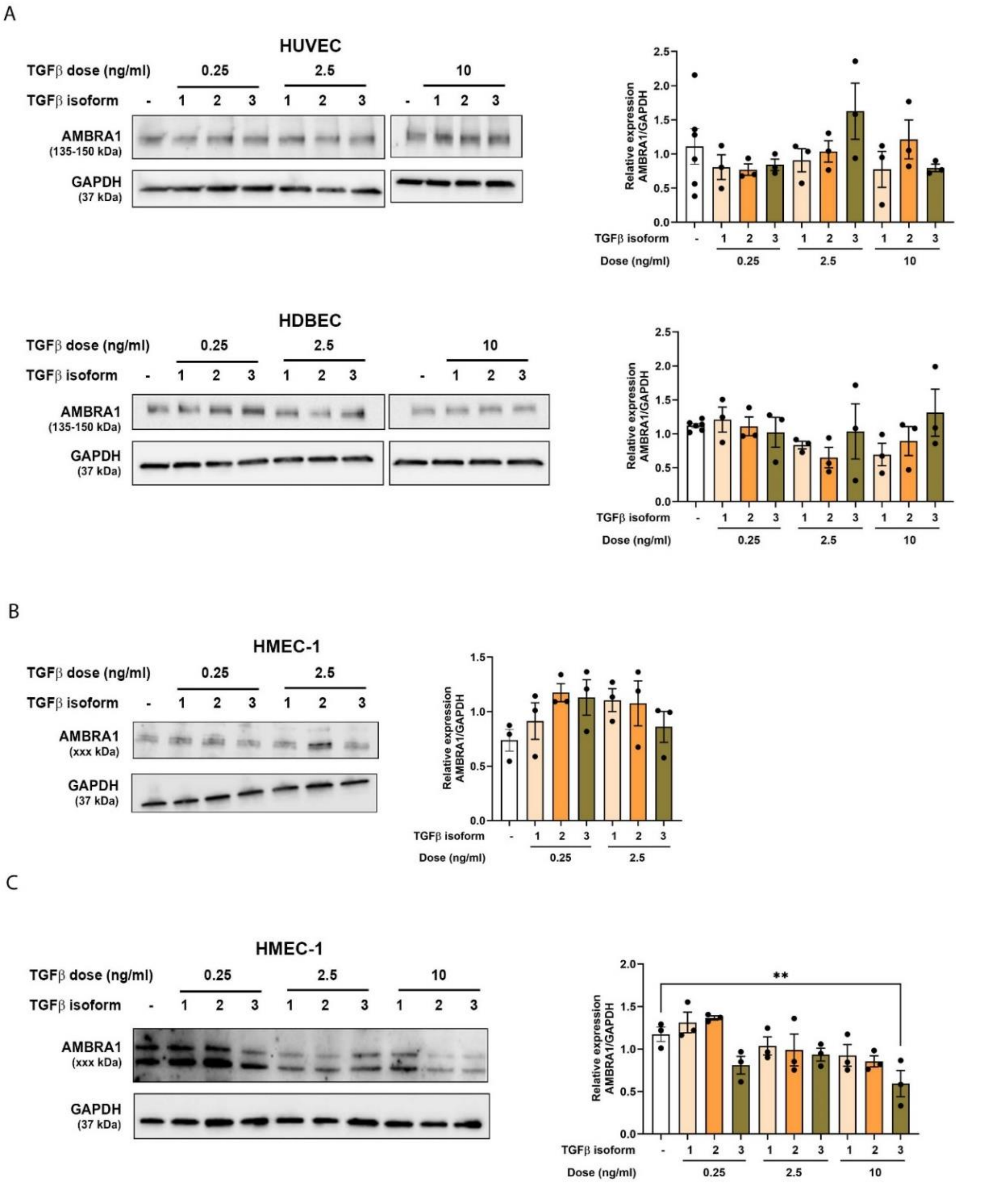

Supplementary Figure 2

A

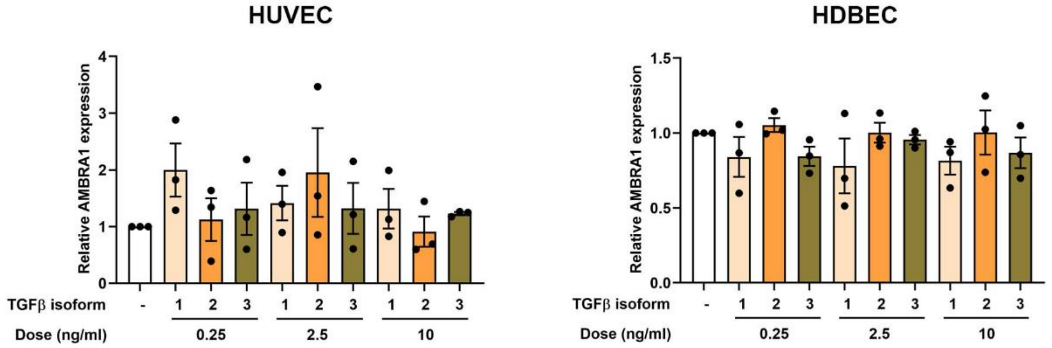


B

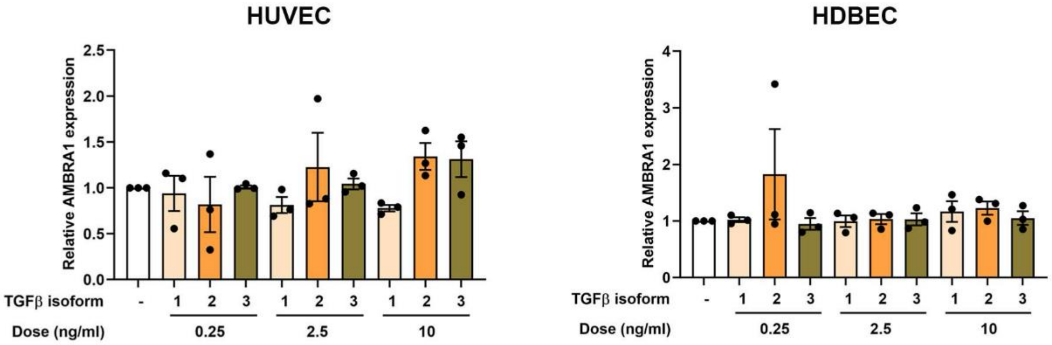

Supplementary Figure 3

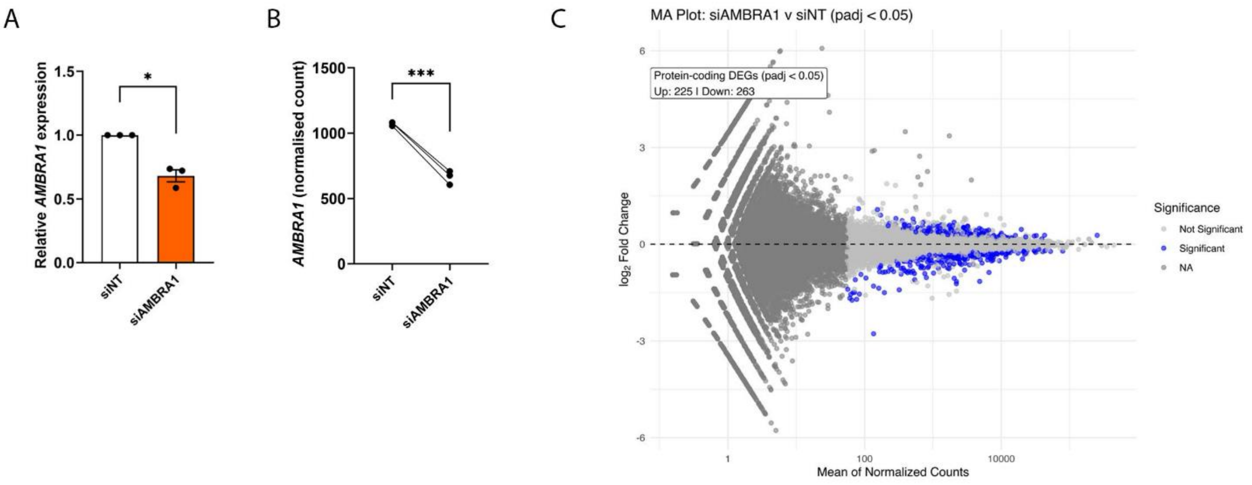

Supplementary Figure 4

A

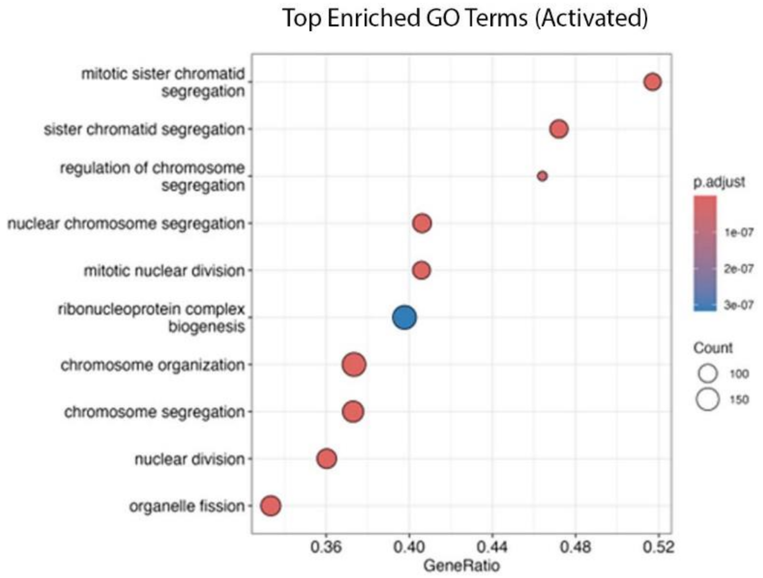


B

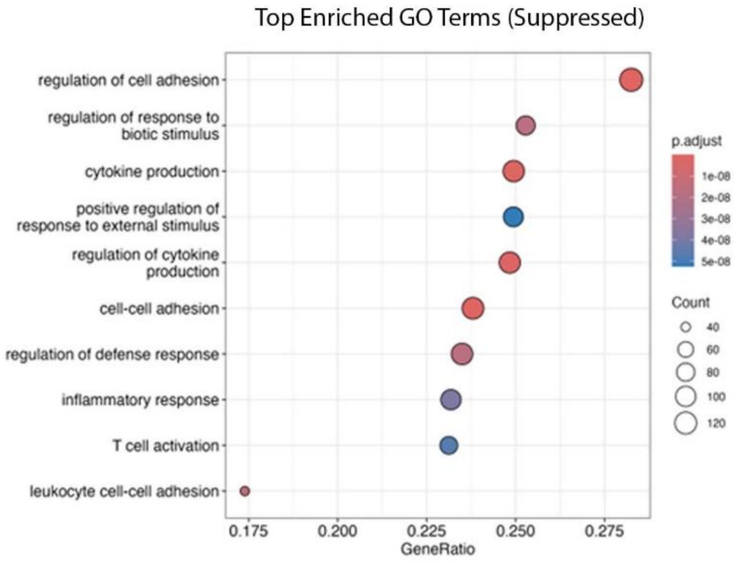

Supplementary Figure 5

A

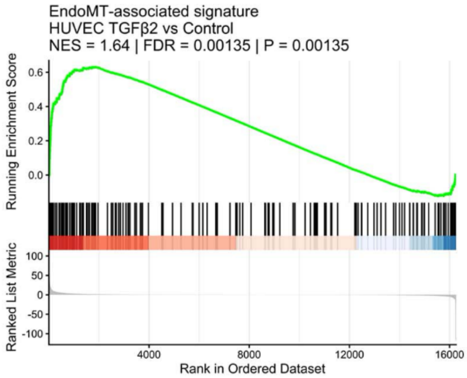


B

| Mesenchymal / mesenchymal-like markers | | |
|---|---|---|
| COL4A2 | CDH2 | LAMC2 |
| FAP | TAGLN | MYL9 |
| COL4A1 | FN1 | CNN1 |
| TNC | COL1A1 | SPARC |
| MMP2 | CALD1 | CNN2 |
| SERPINE1 | ITGB1 | PDGFRB |
| CD44 | POSTN | |
| MMP14 | PDGFRA | |

| EndoMT inducers & signaling receptors | | |
|---|---|---|
| IL1A | TGFB1 | BMPR1B |
| TGFBR1 | BMPR1A | |
| TGFB2 | FGF2 | |
| ACVRL1 | WNT5A | |

| EndoMT positive regulators | | |
|---|---|---|
| KLF4 | NFKB1 | FOXC2 |
| RELB | SNAI1 | RELA |
| NFKB2 | ZEB1 | HEY1 |
| EGR2 | JUN | |

Supplementary Figure 6

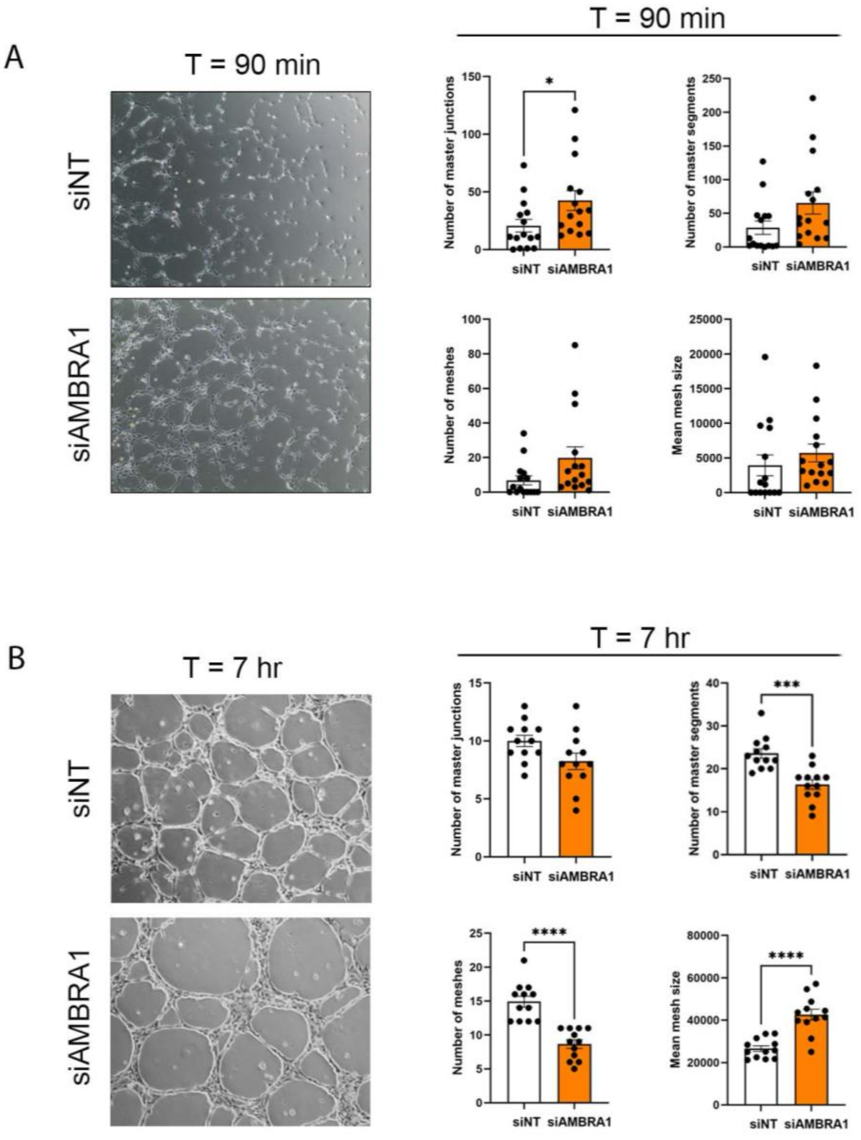

**Figure S1. TGFβ-mediated signalling does not decrease endothelial AMBRA1 levels at 24 hrs.** (A) Western blots of AMBRA1 and GAPDH protein from HUVEC and HDBEC cells in the absence or presence of treatment with TGFβ1, 2 or 3 (0.25-10 ng/ml) for 24 hrs, or from HMEC-1 cells for (B) 24 hrs (0.25-2.5 ng/ml) or (C) 72 hrs. Protein levels were quantified by densitometry, normalised to GAPDH and presented relative to the mean protein/GAPDH value for each experiment (mean ± SEM, *n* = 3) (one-way ANOVA with Dunnett's multiple comparison test on log FC data to compare treatment to control; ** $P < 0.01$).

**Figure S2. TGFβ signalling does not decrease AMBRA1 gene expression.** (A,B) RT-qPCR mRNA expression analysis of AMBRA1 from HUVEC and HDBEC endothelial cells in the absence or presence of treatment with TGFβ1, 2 or 3 (0.25-10 ng/ml) for (A) 24 hrs or (B) 72 hrs. mRNA expression levels were normalised to GAPDH and presented relative to mRNA levels in control vehicle-treated cells (mean ± SEM, *n* = 3).

**Figure S3. Validation of siRNA-mediated AMBRA1 knockdown in RNA-sequencing samples and differential expression profiling in HUVEC cells** (A) RT-qPCR mRNA expression analysis of AMBRA1 from HUVEC cells transfected with control (siNT) or AMBRA1 siRNA for 48 hrs. mRNA expression levels were normalised to GAPDH and presented relative to siNT (mean ± SEM, n = 3; one-sample t-test on logFC data; * $P < 0.05$). (B) Expression of AMBRA1 from DESeq2 normalised counts in siAMBRA1-depleted cells versus a siNT control. Points represent individual biological replicates (*** *padj* < 0.05) (C) DEGs were assessed for AMBRA1 depleted versus non-targeting (siNT) control in the HUVEC cell line. Dots represent protein-coding genes up- or downregulated, demonstrating a 1.5-fold change which have a *padj* value < 0.05 (blue).

**Figure S4. Gene Ontology (GO) GSEA of AMBRA1-regulated transcriptional programmes** Gene Ontology (GO) GSEA was performed using ranked DEGs from siRNA-mediated AMBRA1 depleted HUVEC cells versus non-targeting (siNT) control. Dot plots show the top 10 significantly (*padj* < 0.05) enriched GO terms which were (A) activated (NES > 0), (B) or suppressed (NES < 0). Dot size reflects gene set size and colour indicates significance (*padj*).

**Figure S5. Validation of the curated EndoMT signature.** (A) GSEA of LIMMA-ranked gene expression data from TGFβ2-treated HUVECs (GSE118446) using the curated EndoMT gene signature (B) Leading-edge genes contributing to EndoMT positive gene set enrichment.

**Figure S6. AMBRA1 knockdown enhances endothelial cell vascular network formation.** (A,B) HUVEC cells transfected with control (siNT) or AMBRA1 siRNA for 96 hrs (A) or HMEC-1 cells were transfected with control (siNT) or AMBRA1 siRNA #1 for 48 hrs (B) were seeded in Matrigel and tubule formation monitored over time. Images were taken using a 10X objective, and the number of master junctions, master segments and meshes, and mean mesh size, were quantified at 90 min (A) or 7 hr (B) (mean ± SEM, each data point is a technical replicate from a minimum of four independent experiments; *t*-test, * $P < 0.05$, *** $P < 0.001$, **** $P < 0.0001$)
